# Robust quality assessment of cryo-EM maps, tomograms and micrographs by statistics-based local resolution estimation

**DOI:** 10.64898/2026.02.03.703505

**Authors:** David Kartte, Carsten Sachse

**Affiliations:** Ernst-Ruska Centre for Microscopy and Spectroscopy with Electrons, ER-C-3/Structural Biology, Forschungszentrum Jülich, 52425 Jülich, Germany; Department of Biology, Heinrich Heine University, Universitätsstr. 1, 40225 Düsseldorf, Germany

**Keywords:** Cryo-electron microscopy, cryo-electron tomography, map quality, micrograph quality, tomogram quality, resolution, local resolution

## Abstract

Resolution estimation by Fourier shell correlation (FSC) using half data sets is the standard method for map quality assessment in cryo-EM. Currently, the FSC method is largely used for refined cryo-EM maps in the context of single particle cryo-EM or subtomogram averaging. Here, we extended resolution estimation to assess the quality of electron micrographs, tilt-series and tomograms. We developed a robust statistics-based framework, capable of determining local quality estimates in the above cryo-EM data types. We show that the determined quality values on a micrograph and tomogram level can be used as a particle quality criterion to improve averaged 3D reconstructions. Using local quality assessments of tomograms, we were able to characterize tomogram quality dependence on distance inferred by radiation damage of FIB-milled lamella. This robust resolution-based quality assessment approach suitable for multiple cryo-EM data types opens new possibilities for automated quality control and method development in cryo-EM maps as well as tomograms and micrographs.

## Introduction

Electron cryo-microscopy (cryo-EM) has become a highly popular method for structure determination of biological macromolecules over the last decade (Kühlbrandt, 2014). In particular, single-particle cryo-EM of biochemically isolated molecules routinely yields high-resolution structures of 2-4 Å in resolution, allowing for atomic modelling. Alternatively, subtomogram averaging based structure determination has been employed in cases when macromolecules cannot be sufficiently purified or are too heterogeneous. As the corresponding software solutions have become increasingly mature, subtomogram has also yielded near-atomic resolutions in some cases (Burt et al., 2024; Himes and Zhang, 2018; Tegunov et al., 2021; Zivanov et al., 2022). For single-particle as well as subtomogram averaging cryo-EM structures, resolution estimates are presented as the single most important measure for map quality, indicating the reliability of map features suitable for model interpretation.

For the commonly practiced procedure of resolution estimation, the data is split into two half-sets of particles and two half-reconstructions are generated, from which the Fourier Shell Correlation (FSC) is computed (Harauz and van Heel, 1986). To determine the final resolution, the highest-frequency shell surpassing a threshold of 0.143 is determined (Rosenthal and Henderson, 2003). While some aspects of FSC remain under discussion, e.g. the proposition of alternative thresholds due to the effects of sample size (Beckers and Sachse, 2020; Rohou, 2020; van Heel and Schatz, 2005), the FSC measure with a 0.143 cutoff represents the currently most used method used in the EM databank (Kleywegt et al., 2024). When applied to the global cryo-EM structure, the FSC yields a single quality measurement, making the results of cryo-EM experiments comparable.

Experimental cryo-EM maps, however, often do not exhibit equal spatial and isotropic quality across the volume due to preferred particle orientations, alignment inaccuracies and particle flexibility. In these cases, local resolution determination has been shown to be helpful, providing complementary information on different parts of the molecule by repeated local resolution calculations (Cardone et al., 2013; Kucukelbir et al., 2014). In the case of blocres, the FSC is computed in smaller running subvolumes while for ResMap, local sinusoidal features are evaluated, respectively. Multiple implementations of local resolution estimation exist, often relying on either FSC computations (Punjani et al., 2017; Zivanov et al., 2018; Beckers and Sachse, 2020) or local feature evaluation (Vilas et al., 2018) as a starting point for the generation of locally resolution filtered maps. In the presence or absence of an atomic model, a series of local sharpening routines have been proposed to increase feature interpretability (Bharadwaj et al., 2025; Jakobi et al., 2017; Ramírez-Aportela et al., 2020; Terwilliger et al., 2018). Procedurally, for the FSC estimation, the sequence of calculations can also be reversed, conducting correlation measurements in real space instead of Fourier space while obtaining equivalent FSC values (Penczek, 2020). One additional practical item that needs to be considered for local subvolume FSC computations is the window size. While increasing local information is the explicit aim of the local FSC calculations, motivating the usage of smaller window sizes, the subsequently decreasing signal-to-noise ratio is the limiting factor for reliable resolution estimation (Cardone et al., 2013). As a result, in particular the low-resolution shells contain very small numbers of voxels that make signal detection more difficult. Therefore, the window size in local FSC computations represents a fundamental trade-off between spatial localization and signal reliability. Although a combination of permutation and hypothesis testing corrected by false discovery rate control allows for sensitive signal significance evaluation for each resolution shell (Beckers and Sachse, 2020), the choice of the optimal window size remains an important aspect of FSC-based local resolution estimation.

In addition to structure determination of purified proteins by single-particle cryo-EM, the visualization of frozen-hydrated cells using a series of cryo-EM methods is becoming increasingly popular (Lucas, 2023; Nogales and Mahamid, 2024). The workflow of *in situ* cryo-tomography is often complemented by correlative light and electron microscopy (CLEM) in order to identify the regions of interest. Once located, vitrified cells need to be thinned by a dedicated focused ion beam (FIB) scanning electron microscope (SEM) for the final cryo-EM visualization of the lamellae. Often, micrographs or tilt-series merged into 3D tomograms of the cellular lamellae are acquired in order to locate molecular structures at high resolution or organellar ultrastructures (Bäuerlein and Baumeister, 2021). In this way, near-atomic resolution structures of ribosomes within the cellular environment are increasingly becoming available (Xue et al., 2022). Hardware add-ons like a fluorescence microscope integrated into a FIB-SEM (Berkamp et al., 2023) and enhanced image processing reconstruction routines (Eisenstein et al., 2024) are increasing the throughput of the *in situ* workflow. Ribosome structures derived from STA have been used to assess the utility of new developments, e.g. to characterize radiation damage effects for different gases of xenon, argon or oxygen plasma used in FIB milling (Berger et al., 2023).

Here, we present a new, fully automated software program for local resolution estimation in maps, micrographs and tomograms we termed RESOLVE (RESOLution estimation in Variable Environments). First, we show that this approach yields robust and accurate results: It allows reliable measurements in single-particle cryo-EM maps with high locality close to Nyquist. Second, we extend this approach to low signal-to-noise-ratio (SNR) cryo-EM data, to apply resolution estimation to tilt-series, tomograms and 2D micrographs. The methods are validated by selecting an enhanced subset of single particles and subtomograms resulting in higher resolution reconstructions. In this context, RESOLVE is not only a reliable method for local resolution estimation, but it can also be used to assess micrograph and tomogram quality and ultimately improve the corresponding 3D reconstructions.

## Results

### The RESOLVE procedure of local resolution estimation

In order to develop a robust resolution determination tool suitable for multiple data types in cryo-EM, we established the procedure using typical half-maps of single-particle cryo-EM maps, e.g. of the ribosome (EMD-0194). In this procedure, we iterate through each resolution shell (**Figure 1**): (1) we apply the band-pass filter to the half-maps corresponding to the resolution shell and (2) we compute running correlation values for every map voxel, employing cosine similarity (CS) for real-space measurements in analogy to a previously described approach (Penczek, 2020). The CS is defined as

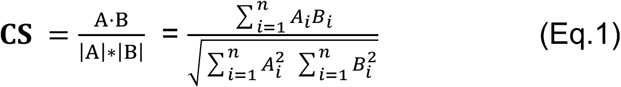

Where *A* and *B* are vectors containing the real-space intensity values within the local window radii of the two half maps, *A·B* is their dot product and *|A|*∗*|B|* the product of their magnitude. This is effectively a real-space equivalent of the FSC:

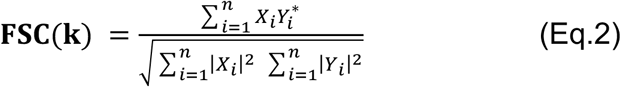

Where *X_i_* and *Y_i_* are factors in Fourier space, thus denoting complex numbers. While for the FSC, only factors within the current shell *k* are included, we use the CS on band-pass-filtered maps, effectively focusing the measurement on a specific shell in Fourier space.

**Figure 1.**
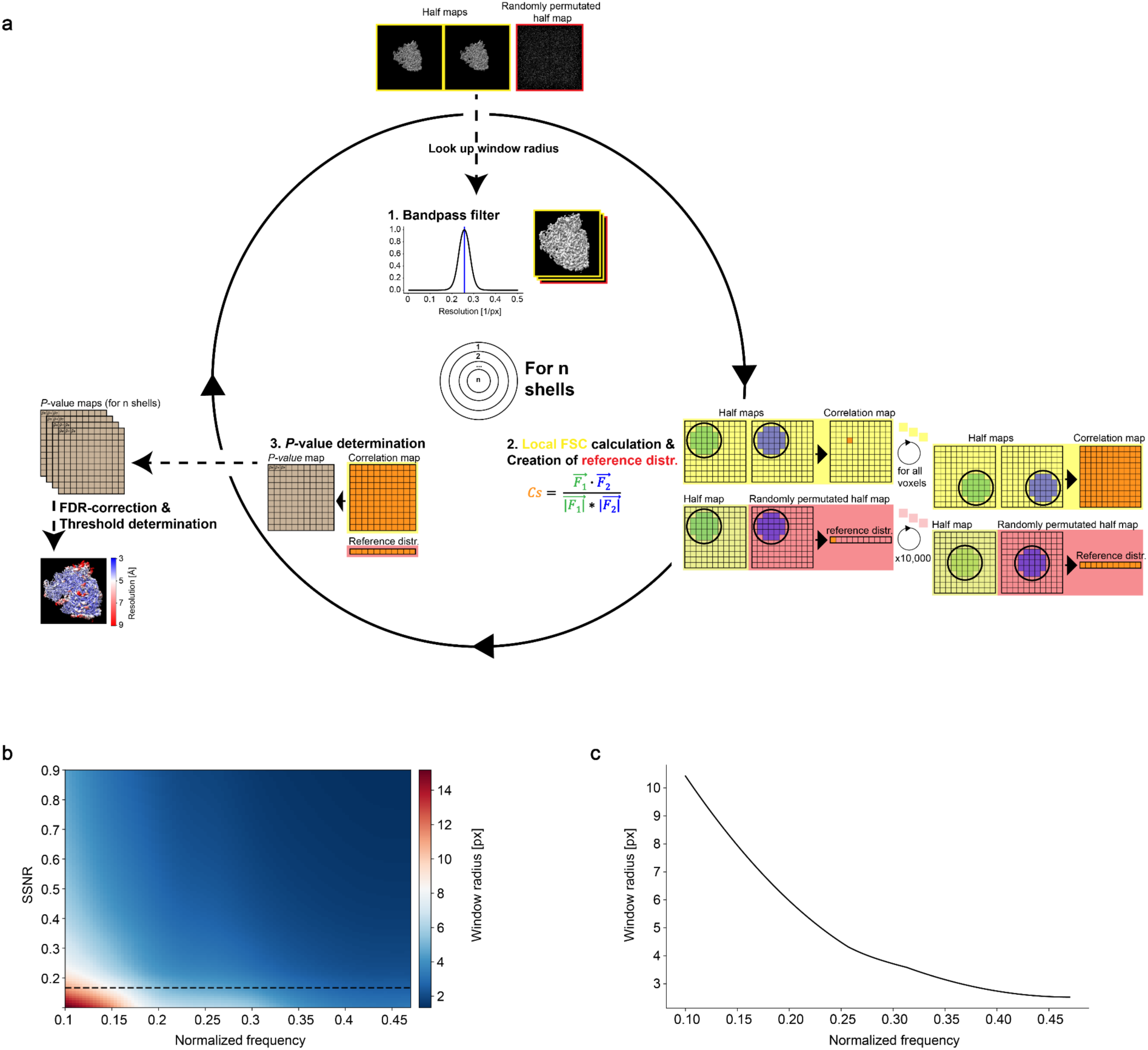
Schematic presentation of the RESOLVE local resolution estimation workflow. a) Steps 1 - 3 are conducted for each resolution shell of the map. (1) Half maps and their permuted counterpart are band-pass-filtered according to the current resolution shell. Resolution-dependent window radii are obtained for each resolution shell. (2) Running correlation calculations are conducted and a reference distribution created. (3) Map of *p*-values (q-values after multiple testing correction) are determined. Finally, after iterating through each shell, FDR-correction is applied and for each voxel of the map a resolution threshold is determined for when the highest shell drops below *q* < 0.01 or 1 %. b) Table with the determined frequency dependent window radii for different spectral signal-to-noise levels (SSNR) (Eq.3), for measurments in 3D maps. SSNR 0.167 (corresponding to a FSC 0.143) is highlighted as a black dotted line. c) Curve with the determined frequency dependent window radii for a SSNR of 0.167, for measurments in 3D maps.

After looking up the predetermined resolution-dependent optimal window radius, the radius is employed to each map measurement location in order to estimate the correlation for a locally masked area. By including a permuted half-map correlation, (3) we obtain statistically relevant *p*-values for each map location. Once steps 1-3 are iterated over every resolution shell, we identify the highest resolution shell with significant correlations for each map location in analogy to previously applied multiplied testing correction (Beckers and Sachse, 2020): for each map location, we analyze the obtained *p*-values for all resolution shells and use multiple testing correction to derive *q*-values from our *p*-values by false-discovery rate control (Benjamini and Hochberg, 1995). Finally, we assign the identified resolution value to each voxel of the cryo-EM map when the *q*-value of the FSC estimate drops below a defined threshold for the first time, e.g. 1 % FDR.

### Consequences of resolution estimation in real space

Instead of excising local volumes and then conducting the common Fourier correlation measurements, we performed the resolution estimation in real space as outlined above. Practically, we generated a series of band-pass filtered half-maps keeping only the information band of the shell of interest (steps 1 and 2). For the implementation of the band-pass filter operation in Fourier space (step 1), we used a hyperbolic tangent to avoid artifacts through spectral leakage. The filter was applied to both half-maps, introducing dependencies that are known to affect the correlation measurements (Beckers and Sachse, 2020; Penczek, 2020). Here, we dealt with these additional dependencies in the following manner (step 2): we created a correlation reference distribution for each resolution shell by permuting one of the original input half-maps, applying the same band-pass filters as for the original half-maps, and conducting correlation measurements between the permuted half-map and the original half-map for 10,000 random locations. As this reference correlation distribution contained the same dependencies as the originally band-pass-filtered half-maps, we were able to derive the correct *p*-value map by determining the rank of each measurement value with respect to the reference distribution. Finally, after running through each resolution shell, we applied multiple testing correction to the *p*-value map to derive the *q*-values for the final per-voxel threshold determination.

### An adaptable window radius for estimating correlation of each resolution shell

A critical parameter for the reliability of the FSC or, more generally, local correlation calculations, is the choice of the real-space window size or radius (Beckers and Sachse, 2020; Cardone et al., 2013; Zivanov et al., 2018). Here, we wanted to automatically identify the radius with the window as small as possible for obtaining resolution values of high locality while still yielding reliable resolution measurements. Therefore, we worked out a strategy to adapt the window radius depending on the investigated spatial frequency. To approximate the optimal radius, we conducted measurements on 1000 simulated half-map pairs with varying spectral signal-to-noise ratios (SSNR) and identified the smallest window radius per shell still capable of reliably detecting the simulated signal (**Figure 1b**, Material and Methods).

### Application and benchmarking of RESOLVE with single-particle cryo-EM maps

To benchmark our proposed approach for local resolution estimation, we compared RESOLVE’s outcomes of the ribosome (EMD-0194) and SARS-CoV-2 spike protein (EMD-34658) maps with commonly used programs: blocres (Cardone et al., 2013), Monores (Vilas et al., 2018), RELION (Zivanov et al., 2018), ResMap (Kucukelbir et al., 2014) and SPOC (Beckers and Sachse, 2020). In comparison, when the local resolutions were mapped on the isosurface, the different programs yield overall similar results in the relative variation of resolution while absolute resolution values differed (**Figure 2**). We then inspected local resolution values obtained from a single map slice cut through the volume. High variations within short distances represent resolution assignments with high locality (e.g. Monores and ResMap), while using a large window leads to a blurred appearance (e.g. RELION). Clearly, excessively large resolution differences between neighboring pixels are the result of unreliable measurements, and in such circumstances, we found high resolution outliers associated in unexpected locations. The central parameter that determines the balance between high locality and information content is the window size, which can be tuned by the operator for most programs whereas RESOLVE is using adaptable and frequency-dependent window radii. When comparing RESOLVE and blocres, at first glance, they yield similar results, especially for low resolution regions. For higher resolutions, however, RESOLVE revealed more locality. The adaptable window size used in RESOLVE can make use of smaller windows while still yielding reliable results in high resolution regions, and for lower resolution regions, it can refer to larger window sizes. In contrast, blocres remains conservative for high-resolution shells, using the same window size required for reliable measurements of the lowest resolution shell, sacrificing locality and therefore available high-resolution information. RESOLVE’s optimized adaptable window sizes for different resolution shells accommodate a good balance between high-resolution information alongside robust low-resolution resolution estimation.

**Figure 2.**
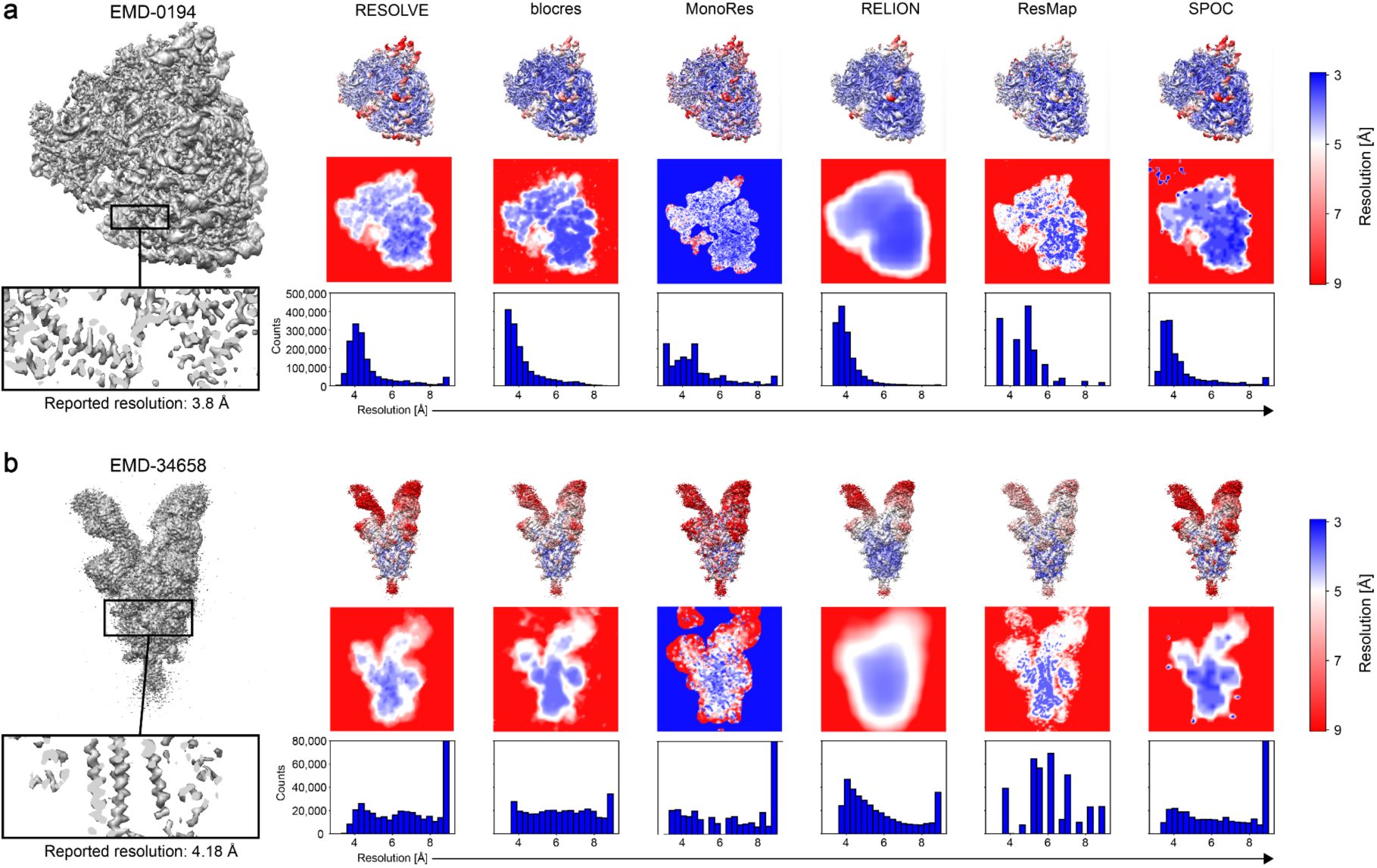
Local resolution estimation for 3D cryo-EM maps. Resolution analysis for a) Rabbit 80S ribosome (EMD-0194) and b) SARS-CoV-2 Omicron BA.1 spike trimer (EMD-34658). For a) and b), the following information is shown in the panels: Top left: cryo-EM map with inset slice through α-helical density. Top right: Results of local resolution estimation using RESOLVE, blocres, MonoRes, RELION, ResMap and SPOC (FDR-FSC). Top row: isosurface with mapped local resolution values (color bar right). Center row: slice through volume center with color density display according to local resolution. Bottom row: Histogram of local resolution values for each program above.

### Computing the reference distribution for different data types

In order to extend the RESOLVE approach to other EM data types, we generated the reference distribution for averaged data from refined maps differently than data derived from single recordings, namely micrographs, tilt-series and tomograms. Refined maps derived from single-particle cryo-EM and subtomogram averaging contain continuous information bands from low to the highest resolution. For these data derived from refined maps, we created permuted half-map by random permutation in real space (Beckers and Sachse, 2020), destroying any signal and keeping only the intensity distribution intact. However, data derived from a single recording portray measurement-specific signal-to-noise characteristics, e.g., according to the contrast transfer function, which are necessary to be retained in the reference distribution. Keeping the signal characteristics intact is equally relevant for tomograms: The reconstruction in Fourier space and the missing wedge introduce additional artificial dependencies represented as consistent characteristics in the power spectrum. To include these effects, we used phase-permutation in Fourier space rather than real-space permutation, keeping the power spectrum of the permuted map intact. To demonstrate that this procedure is capable to compensate for tomographic reconstruction (**Suppl. Figure 1a**) and missing wedge artifacts, we created a test case by permuting every tilt of one half-tilt-series individually, reconstructing the tomogram, to then use it as permuted half-tomogram for reference distribution creation, leading to a realistic null distribution. Indeed, for this test case, we measured slightly more conservative resolutions when compared with the reference distribution by simple real-space permutation (**Suppl. Figure 1b-c**), indicating that artificial dependencies introduced by the tomogram reconstruction procedure inflate correlation measurements by small margins. Moreover, we employed phase-permutation instead of real-space permutation on the half-tomograms also preventing the artificial inflation of resolution measurements (**Suppl. Figure 1d**), as it keeps the permuted power spectrum with is systematic characteristics intact. Therefore, for non-averaged measurements with a discontinuous information spectrum like micrographs, tilt-series and tomograms, from here on, we compute the reference distribution by phase-permutation.

### Local resolution estimation of two-dimensional (2D) micrograph movies

After establishing the RESOLVE work-flow for three-dimensional (3D) cryo-EM maps, we set out to apply it to micrograph movies to determine local resolution estimates and derive a global micrograph quality criterion. Here, we computed the 2D equivalent in real space using the Fourier ring correlation, obtained from half-sets made of even and odd frames of each micrograph (**Figure 3a**). The generation of the lookup table for the optimal window radius was also adjusted to account for the smaller number of pixels involved (Materials and Methods, **Suppl. Figure 2a, b**). The remaining workflow steps are identical to the way described above for 3D maps. First, we tested the method on a previously recorded TMV micrograph ‘TMV_004_Sep18’ (EMPIAR-10305). The local resolution mapped on the motion-corrected micrographs showed resolutions up to 6 Å in areas of TMV as opposed to empty areas of vitreous ice between 20 and 30 Å. When we compared the motion-corrected sum of the aligned frames versus the simple sum of the unaligned frames, we detected a significant deterioration in resolution in areas of TMV and vitreous ice (**Figure 3b**). For single micrographs, one should however keep in mind that a single high-resolution measurement cannot be fully representative of the information content due to the commonly applied underfocus in cryo-EM image acquisition. To investigate the effect of the CTF, we plotted frequency-dependent changes of the median *q*-values obtained from the entire micrograph against the estimated CTF by CTFFIND4 (Rohou and Grigorieff 2015), showing a clear inverse correlation, i.e. the *q-*values are lowest in the regions of high contrast transfer, whereas they are high in the regions of the zero intercepts. To mitigate this effect for the case of micrograph measurements (as well as tilt-series or tomograms below), we specifically chose to adjust our final threshold determination step, using a *q*-value cutoff of 0.05 instead of 0.01. In this context, we also propose a single global resolution estimation based on the local estimates of the micrograph. We term this measurement median resolution, as we derive it from a list of shell-associated median *p*-values (details in Material and Methods).

**Figure 3.**
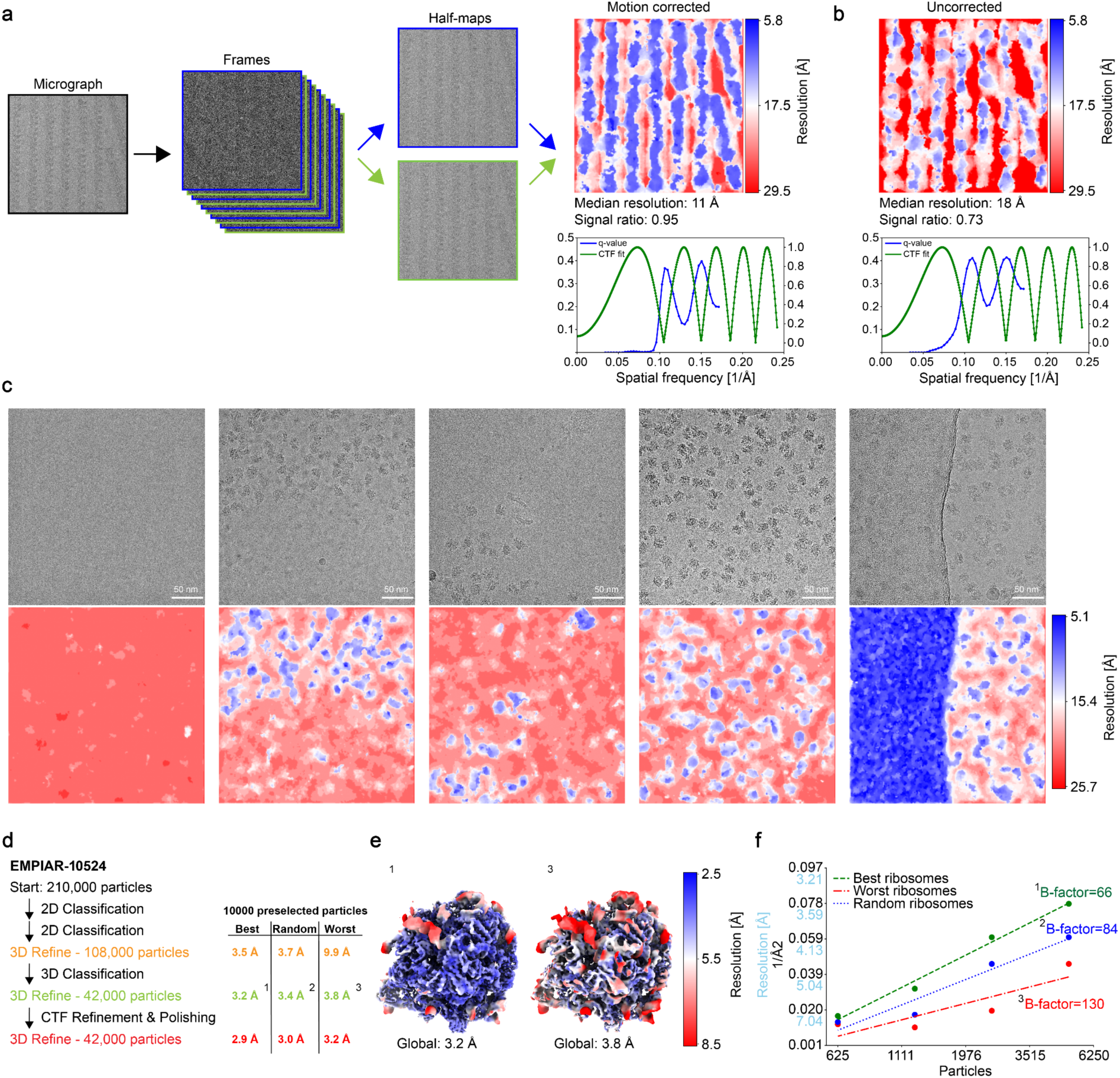
Application of RESOLVE to 2D electron micrographs. a) Micrograph frames are split into odd and even half-sets. Local resolution map of a motion corrected micrograph from tobacco mosaic virus (TMV) (EMPIAR-10305). Below the micrograph: plot of the median *q*-values from our measurements compared with a CTF-fit of CTFFIND4 for the same micrograph. b) In contrast, resolution estimation of a simple sum of aligned TMV micrograph frames (not motion-corrected). Below the micrograph: plot of the median *q*-values from our measurements compared with a CTF-fit of CTFFIND4. c) A set of micrographs (top) with their resolution estimates (bottom) of E. coli 50S ribosomes (EMPIAR-10524). The columns show from left to right: an empty micrograph, a density gradient, a mostly empty micrograph with a strong signal from an artifact in the top right corner, a regular micrograph and a micrograph partially containing carbon film. d) Left: Basic procedure of single-particle workflow (EMPIAR-10524). Right: Global resolution estimates for 10,000 best, random and worst preselected particles by RESOLVE on a micrograph level. e) Local resolution as estimated by RESOLVE mapped on 3D refinement results of best and worst particles, locally filtered maps. f) B-factor comparison for ‘best’, ‘random’ and ‘worst’ particles as determined by RESOLVE.

When RESOLVE estimated the local resolution of micrographs from E. coli 50S ribosomes (EMPIAR-10524), we were also able to confirm higher resolution estimates for areas with particles present and lower in case they were absent (**Figure 3c**). However, we also noticed two additional cases leading to high resolution: first, in the case of devitrified ice and second when carbon film was present. While operating with the median resolution, those effects can be minimized depending on the extent present in the micrographs. For the purpose of the following tests, we excluded those micrographs including carbon film. After showing the principal utility of RESOLVE for estimating local resolution on micrographs, we set out to apply it as a quality criterion for imaged particles on a micrograph level. We reasoned that when better resolved single particles should give rise to improved 3D maps. For that purpose, we reprocessed a ribosome dataset (EMPIAR-10524) and selected the best and worst 10,000 particles as determined by RESOLVE on a micrograph level. We processed these two subsets ‘best’ and ‘worst’, in addition to a ‘random’ subset with randomly chosen 10,000 particles. In order assess the effect of image processing, we conducted this procedure at three different points of the processing workflow (**Figure 3d**): First, after two rounds of 2D classification, second, after one further round of 3D classification and third, we subjected this set of particles to a round of CTF-refinements and particle polishing following the RELION documentation. For those three types of refinements, we obtained consistently highest resolution when using the ‘best’ particle set when compared with the ‘random’ or ‘worst’, where the lowest resolution was obtained. For instance, after 3D classification with a total of 42,000 particles, RELION’s resolution estimates (FSC 0.143) of the cryo-EM map were 3.2, 3.4 and 3.8 Å for the ‘best’, ‘random’ and ‘worst’ particle sets, respectively. Visual comparison of the ‘best’ vs. ‘worst’ maps confirms the resolution estimate (**Figure 3e**). The fitted Rosenthal-Henderson B-factors at 66, 84 and 130 Å^2^ for ‘best’, ‘random’ and ‘worst’ particle sets, respectively, further supports the notion that preselection based on RESOLVE’s local resolution can potentially be used to improve the 3D map quality (**Figure 3f**). Together, RESOLVE is capable of estimating the local resolution on micrograph movies. The obtained global and local quality criteria can be used to select for better subsets of micrographs and particles, resulting in improved cryo-EM reconstructions.

### Application to tilt-series and tomograms

After successfully applying our local resolution estimation tool RESOLVE to 2D micrograph movies, we investigated its utility for cryo-electron tomography. First, we sought to evaluate RESOLVE for tilt-series quality measurements and later on for entire tomographic reconstructions. For the tilt-series acquired on a lamella from -61 to 40°, we applied RESOLVE to each tilt micrograph based on the half-set frames, estimating local resolution in analogy to 2D micrographs (**Figure 4a, Suppl. Figure 3**). For a more compact visualization, we displayed the median *p-*values for each spatial frequency and stacked them vertically for each tilt resulting in a heat map. The resulting map has a pyramid-like shape, where the lowest tilts show highest resolution. This finding can be rationalized as the data was acquired dose symmetrically, i.e. the planar tilts were recorded first in order to minimize radiation damage. Therefore, likely due to radiation damage and sample thickness, tilts that were recorded later showed consistently poorer resolution estimates, giving rise to the pyramid shape of the heat map. As expected, the above-described effects of the CTF could be observed as well, as *p*-values inversely correlate with the defocus and drop almost to 0.01 for higher frequencies a second time. RESOLVE’s local resolution estimates capture the deterioration of resolution for tilted micrograph acquisitions due to buildup of radiation damage and higher tilt angles.

**Figure 4.**
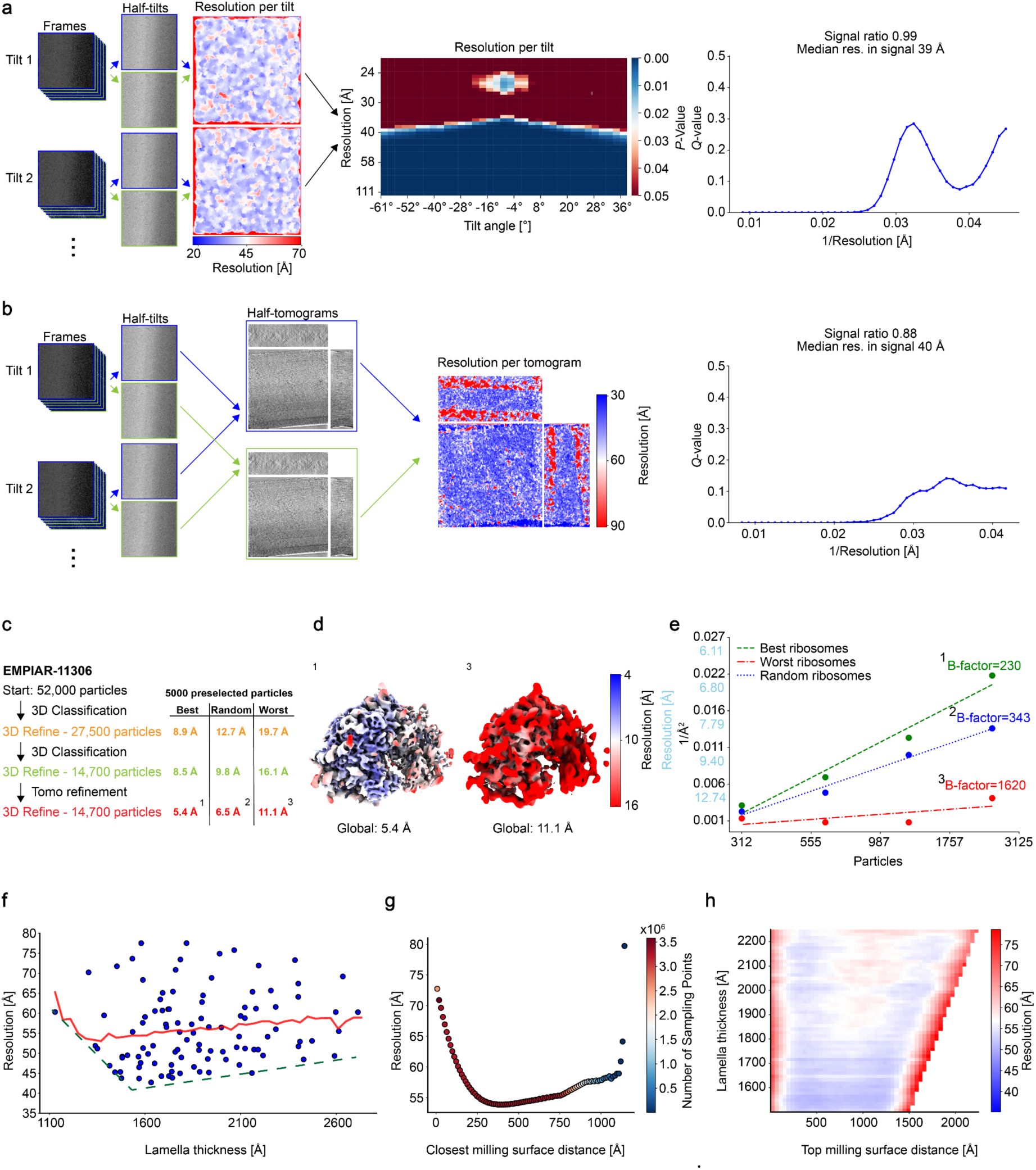
Application of RESOLVE to cryo-electron tilt-series, tomograms and subtomograms. a) Procedure of dividing a tomographic tilt-series into two half-tilt-series. Middle: resolution measurements displayed as heat map of vertically stacked tilts with median *p*-value per resolution. Right: median *q*-values per resolution. b) Procedure for dividing a tomogram into half-tomograms. Middle: local resolution measurements displayed in tomograms. Right: median *q*-values per resolution. c) Top: Basic workflow of subtomogram averaging (EMPIAR-11306). Bottom: RELION global resolution estimates for 5000 ‘best’, ‘random’ and ‘worst’ preselected ribosome subtomogram based on RESOLVE’s local resolution assessment. d) Local resolution as estimated by RESOLVE mapped on 3D refinement results of ‘best’ and ‘worst’ particles, locally filtered maps. e) B-factor comparison for ‘best’, ‘random’ and ‘worst’ particles as determined by RESOLVE. f) Scatter plot of average resolution per tomogram (within lamella) vs. lamella thickness (n=106). Low resolution outliers >= 80 Å were removed (plot with full data in Suppl. Figure 6a). Moving average as red line (window size 400 Å), green dotted line corresponds to high resolution limit. g) Scatter plot of average resolution vs. distance to closest milling surface (within lamella). Only data points from tomograms with average thickness of 150 nm - 225 nm are shown (plot with full data in Suppl. Figure 6c). h) Lamella thickness vs. distance to top milling surface, resolution is color coded. Only data points from tomograms with average thickness of 150 nm - 225 nm (plot with full data in Suppl. Figure 6e).

Next, we applied RESOLVE for 3D local resolution estimations of reconstructed 3D electron tomograms (**Figure 4b**). In some cases of our investigated dataset, when the tomogram quality was poor due to the lack of clear features during tracking, the quality of the tomogram as judged by visual inspection coincided with RESOLVE’s resolution estimates when the resolution was worse than 90 Å (**Suppl. Figure 4a**). However, in many cases, it was not easily possible to judge tomogram quality by visual inspection only. In such cases, RESOLVE’s resolution estimates provided good guidance for tomogram quality assessment up to 30 Å resolution (**Suppl. Figure 4b-d**). To further validate our resolution estimates, we turned to another benchmarking approach involving the local quality assessment of particles located in subtomograms, analogous to the presented cases on single-particle cryo-EM. Using ribosomes from the dataset EMPIAR-11306, initially we started with a total of 52,000 particles, and generated a cleaner subset of 14,700 particles due to heterogeneity by 3D classification (**Figure 4c**). Next, we used RESOLVE to assess local tomogram quality, again creating three subtomogram subsets of 5000 particles, i.e. according to the ‘best’, ‘random’ and ‘worst’ local resolution criteria. We performed subtomogram averaging before running the RELION tomo refinement cycle yielding 8.5, 9.8 and 16.1 Å resolution (FSC(0.143) criterion RELION), and after running the RELION tomo refinement cycle yielding 5.4, 6.5, 11.1 Å for ‘best’, ‘random’ and ‘worst’ particles, respectively. Surprisingly, large differences in resolution and in map quality could be detected between best and worst particles for all three stages of refinement (**Figure 4d**). The Rosenthal-Henderson plot and the fitted B-factors of 230, 343 and 1620 Å^2^ (best, random, worst, respectively) (**Figure 4e**) confirmed the previous single-particle observations that the preselection based on RESOLVE’s tomogram local resolution estimates presents an alternative way to assess particle quality for larger data sets at an early stage.

As RESOLVE was able to demonstrate robust local tomogram quality assessment useful for subtomogram averaging, we wanted to investigate other matters related to resolution in cryo-tomography. In order to examine the effects of damage inferred by the high-energy beam of a focused ion beam (FIB) milling instrument, we further investigated EMPIAR-11306. First, we investigated single tomograms, only focusing on the ribosome coordinates as used for subtomogram averaging by RELION. We determined the local tomogram resolution with RESOLVE and averaged the values within 20 nm boxes to estimate resolution at the ribosome location. When those resolutions are mapped as a function of lamella z-position, the best resolutions 35-40 Å can be observed in the center of the lamellae whereas the resolution drops to 60-70 Å close to the lamella surface (**Suppl. Figure 5a**). This effect is more pronounced for thin lamellae than for thicker lamellae (**Suppl. Figure 5b**). RESOLVE’s local resolution analysis of lamellae is in line with previous observations of resolution deterioration towards the lamellae surface caused by radiation damage (Berger et al., 2023; Lucas and Grigorieff, 2023).

For further investigation of tomogram quality, we quantified RESOLVE’s local resolution estimates across the full tomographic dataset. First, we plotted the average resolution of voxels inside FIB-milled lamellae per tomogram as determined by RESOLVE dependent on corresponding lamella thickness (**Figure 4f, Suppl. Figure 6a** for full data). While the relationship may not be obvious at first glance, averaging resolution estimates reveals that high lamella thickness coincides with poorer resolution on average (**Figure 4f, red line**). This observation is in accordance with the high-resolution limit that equally relates to lamella thickness (**Figure 4f, dotted green line**). Best resolved tomograms were found in lamellae around 160 nm thickness, while for thinner lamellae the resolution becomes poorer again. As diverse tomogram content contributes to differences in resolution, we compared the overall mean resolution of voxels included in a lamella with the mean resolution focused on the position of ribosomes, i.e. within 20 nm boxes around ribosome coordinates (**Suppl. Figure 6a-b**) supporting a similar trend of highest resolution observed for around 160 nm thick lamella.

Next, we investigated the resolution distribution within lamellae and plotted the average resolution against the distance to the closest lamella surface (top or bottom) across the dataset (**Figure 4g, Suppl. Figure 6c for full data**). When the distance from the closest surface increased, we observed an improvement in resolution from 70 Å until it reached a resolution plateau around 50 Å between 35-60 nm distance. Beyond 60 nm distance, the resolution became worse again at a slower rate, which is probably due to the contribution of thicker lamellae to the data set. In addition, we also analyzed the voxels around ribosome positions across the full dataset, yielding a very similar distribution (**Suppl. Figure 6d**). Due to the large amount of data for a range of lamella thickness between 100 and 300 nm, we sorted and stacked the resolution distribution as a function of increasing lamella thickness (**Figure 4f, Suppl. Figure 6e for full data**). Interestingly, for 150 – 180 nm thick lamella, the best resolutions are found in the center of the lamella whereas for lamella above 180 nm thickness, quality appears to decrease in the lamella center with respect to adjacent areas. Together, RESOLVE is capable of an in-depth resolution characterization of FIB lamella and in this context it is able capture resolution deterioration induced by the radiation damage of focused ion milling.

## Discussion

So far, local resolution estimation has been mostly applied as a tool for measuring quality of reconstructed cryo-EM maps and has become a routine representation in many publications (Beckers and Sachse, 2020; Cardone et al., 2013; Kucukelbir et al., 2014; Vilas et al., 2018). In this manuscript, we introduce RESOLVE that is suitable for a much larger set of EM data types such as micrographs, tilt series, tomograms as well as averaged cryo-EM maps. Our tool provides a robust local resolution estimation approach based on an optimized tradeoff between measurement accuracy and measurement locality per resolution shell (**Figure 1**). We applied and benchmarked RESOLVE for local resolution estimation beyond averaged maps, applying it to micrographs, tilt-series and tomograms (**Figure 2-4**). Given the robustness, we found RESOLVE to be beneficial for automated quality assessment of micrographs and tomograms, pre-selecting particles on micrographs, pre-selecting subtomograms within tomograms even revealing resolution deterioration effects like radiation damage effects through FIB milling.

Given the demonstrated utility in applying the same basic approach to so many different EM data types, we believe that the combination of choosing the optimal window radius with the statistical thresholding method to be the key for local and reliable measurements. The proposed approach is different to most other available programs that rely on the user-defined tuning of a fixed FSC cutoff and on a fixed window size for a given SNR (Cardone et al., 2013; Punjani et al., 2017; Zivanov et al., 2018). Choosing window radii dependent on spatial frequency instead of SSNR has the advantage of using comparable measurement conditions and statistics for each measured frequency, as a reliable measurement requires a minimum number of frequencies present in the local windows as well as sufficient sampling of the resolution shells. Due to the window radii being independent of the SSNR of the input half-maps, this strategy also avoids the difficulties in comparing resolution measurements derived from different window sizes among maps with differing SSNRs. RESOLVE runs on a typical workstation including GPU for large maps, typically taking less than 20 seconds for micrographs and up to several minutes for larger tomograms (**Table 1**). For a single-particle cryo-EM map of 170 MB size, RESOLVE completed processing in 1:50 min requiring 5.5 GB of RAM and 1.5 GB of GPU memory. For single micrographs of 2.7 MB size, processing took only 12 s while for standard 0.56 GB tomograms 4:06 min.

**Table 1:**

| Number of pixels/voxels | RAM mem (GB) | GPU mem (GB) | GPU specification | Number of GPUs | CPU specification | Runtime (min:s) |
| --- | --- | --- | --- | --- | --- | --- |
| Micrographs |  |  |  |  |  |  |
| 824 x 852 | <0.1 | - | - | - | AMD Ryzen 9 7900X 12-Core | 0:12 |
| 824 x 852 | <0.1 | - | - | - | AMD Ryzen 7 7700X 8-Core | 0:13 |
| Cryo-EM maps |  |  |  |  |  |  |
| 356 x 356 x 356 | 5.5 | 1 | NVIDIA GeForce RTX 3060 | 2 | AMD Ryzen 9 7900X 12-Core | 1:36 |
| 356 x 356 x 356 | 5.5 | 1 | NVIDIA GeForce RTX 3060 | 1 | AMD Ryzen 9 7900X 12-Core | 1:48 |
| 356 x 356 x 356 | 5.5 | 1.4 | NVIDIA GeForce RTX 3060 | 2 | AMD Ryzen 7 7700X 8-Core | 2:11 |
| Tomograms |  |  |  |  |  |  |
| 694 x 694 x 308 | 14 | 3 | NVIDIA GeForce RTX 3060 | 2 | AMD Ryzen 9 7900X 12-Core | 4:06 |
| 694 x 694 x 308 | 14 | 3 | NVIDIA GeForce RTX 3060 | 1 | AMD Ryzen 9 7900X 12-Core | 4:50 |
| 694 x 694 x 308 | 14 | 3 | NVIDIA GeForce RTX 3060 | 2 | AMD Ryzen 7 7700X 8-Core | 6:16 |

We developed a general framework for the characterization of multiple EM data types such as cryo-EM maps, micrographs and tomograms. While global resolution can be used as an overall metric, local quality metrics are better suited to assess local properties for a given data type. Quality assessment based on global resolution metrics such as CTF fit and resolution measures of class averages are routinely performed in cryo-EM data processing (Punjani et al., 2017; Rohou and Grigorieff, 2015; Zivanov et al., 2018). To make a convincing case for local resolution metrics on micrographs and tomograms, we selected particle quality based on RESOLVE’s local resolution estimates and confirmed improvements for single-particle cryo-EM and subtomogram averaging. Additional results indicate that RESOLVE works particularly well to estimate local quality for *in situ* lamella tomograms. Derived from RESOLVE’s local resolution measurements, we also provide a median global resolution estimate, which closely corresponds to the estimated global resolutions. Robust local and global quality assessments will prevent data overinterpretation and can be used for early data cleaning: either for particle curation for structure determination or for automated tomogram analysis, facilitating faster and more accurate downstream processing. As tomography workflows become increasingly automated, RESOLVE may thus provide an alternative approach of assessing data quality directly after reconstruction.

In this manuscript, we showed that RESOLVE can facilitate micrograph, tilt-series and tomogram quality characterization. For instance, we show that for micrographs, RESOLVE can assess the effect of motion correction. For measurements of 2D tilt-series, RESOLVE adequately captures quality differences between earlier and later recorded tilts. This finding indicates that RESOLVE’s measurements account for the effects of radiation damage on data quality (Baker and Rubinstein, 2010). For tomograms, we used RESOLVE for the detailed characterization of quality dependence on distance from lamella surface in accordance with other previous thickness characterizations (Tuijtel et al., 2024). Such local quality studies for *in situ* tomograms have previously been conducted by computationally expensive subtomogram averaging (Baker and Rubinstein, 2010; Berger et al., 2023) or 2D template matching (Lucas and Grigorieff, 2023). Our results indicate that tomogram quality is poor in the area within ∼35 nm distance of the lamella surface and improves toward the lamella center. However, for lamella thicker than 175 nm we also observed a deterioration of quality in the lamella center. With the given tomographic imaging and reconstructions approaches, lamellae of 160 nm appear to yield best resolution values. In this context, we believe that RESOLVE will become useful for assessing different data acquisition strategies and image processing routines.

### Limitations

Given the demonstrated cases of resolution assessments, it is worthwhile to mention that RESOLVE will be equally mislead by averaged cryo-EM maps generated through overfitting procedures during refinement to report overoptimistic resolution (Grigorieff, 2000). RESOLVE is based FSC measurements and, therefore, other precautions need to be taken to avoid overfitting (Chen et al., 2013; Scheres and Chen, 2012), which are typically implemented in the current most popular single-particle workflows (Punjani et al., 2017; Scheres, 2012). In addition, there is one important consideration for local resolution estimation as used with the proposed cryo-EM data types: although RESOLVE effectively characterizes the SNR, the result is not necessarily related to quality of the signal of interest. Averaged maps are *de facto* generated using the signal of interest, as many selected particle images are used to enhance the underlying signal. In the averaging context, any other object or processing artifact that itself has high SNR, is neglected or suppressed in the procedure. However, when the data are not averaged, high SNR regions arising from strong electron scattering or imaging artifacts will be assigned high resolution to and are in that moment impossible to differentiate from the signal of interest. We demonstrated this effect for the following examples: carbon film (**Fig. 3**), devitrified ice (**Fig. 3., Suppl. Figure 4**), reconstruction artifacts at tomogram surfaces (**Suppl. Figure 4**), or sometimes regions in tomograms far outside the lamella (**Suppl. Figure 4**). For the assessment of local features, those cases can be quickly identified by eye and flagged when the signal is magnitudes stronger than the rest of the data. For the judgement of quality by global metrics using median local resolution, this effect is less problematic as the high SNR regions are of limited dimension and they have little influence on the median global resolution and will not affect global quality assessment. For larger scale studies, quantification over many tomograms make such effects more negligible. One additional limitation should be noted that in the context of single-particle cryo-EM or subtomogram averaging, RESOLVE’s resolution estimates do not necessarily capture structural homogeneity, which is one important factor constituting map quality in the context of structure determination. Thus, RESOLVE’s function should be understood as a complementary tool for particle selection rather than an alternative to conventional classification algorithms.

## Acknowledgments

The authors acknowledge support of the project KA1-Co-02 ‘CoViPa’, a grant from the Helmholtz Association’s Initiative and Network Fund. The authors gratefully acknowledge the computing time granted through Jülich Aachen Research Alliance (JARA) on the supercomputer JURECA at Forschungszentrum Jülich .

## Author contributions statement

D.K. and C.S. conceived the work. D.K. was involved in programming the RESOLVE.

D.K. and C.S. were analyzing data and in writing the manuscript.

## Methods

### Estimation of optimal window radii for FSC measurements

In order to determine the optimal window radii for the FSC measurement, we relied on simulating defined SSNR with synthetic data. For our simulations, we chose the atomic model of human parechovirus (PDB-ID 4udf) that exhibited a high surface-to-volume ratio area and unevenly distributed signal of a hollow sphere across the map. For each resolution shell we tested, we created a new set of simulated maps with systematically varying SSNR. More specifically, we sampled the frequency domain for 10 resolution shells, covering the frequency spectrum evenly with the lowest at a normalized frequency of 0.1 and the highest at 0.47 close to the Nyquist limit. For each of these 10 resolution shells, we simulated 100 different half-map pairs of varying SSNRs ranging from approximately 0.9 to 4.4 by scaling the mean of the white noise power spectrum relative to the mean of the shell-specific power spectrum of each simulated map. For each shell-specific simulated map with known SSNR, we now performed the RESOLVE algorithm for local correlation estimation only focusing on the map-specific shell, using a series of differing window radii ranging from 2 to 30 pixels for 3D. We defined correct signal detection as 95% of signal regions derived from the noise-free maps passing the *p*<0.01 threshold when testing correlation against the reference distribution. This way, we identified the smallest window still detecting the known SSNR correctly. We linearly interpolated between our SSNR/FSC specific sampling points to create an extensive lookup table (**Suppl. Figure 2a, left**) and extracted the smoothened curve for the relevant SSNR value of 0.167. Given that SSNR is related to the Fourier shell correlation value at a given resolution shell:

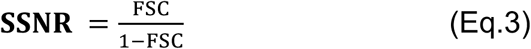

the simulated SSNR of 0.167 corresponded to practically relevant FSC value of ∼0.143, allowing the calculated windows to be large enough to reliably identify signal up to that noise level. To smoothen the derived window radii relationship with spatial frequency (**Suppl. Figure 2a, middle**), we used a Savitzky-Golay filter (Savitzky and Golay, 1964) (**Suppl. Figure 2a, right**). In this way, using simulations of known SSNRs, we generated a lookup table with the smallest appropriate window size to still reliably estimate resolution at the corresponding spatial frequency. We repeated the same procedure for 2D maps (**Suppl. Figure 2b**).

### Identification of window radii for tomograms, tilt-series and micrographs

Due to the missing third dimension, larger window radii are required to include sufficient number of pixels for reliable measurements. However, in particular for lower frequencies, using larger window radii comes at the expense of the locality of the measurements. Consequently, the required window radii for reliable correlation calculations are higher in 2D, and in our simulation, we tested radii between 2 and 100 pixels. Therefore, for the window radii assessment in tilt-series and micrographs, we adjusted the parameters during the simulation to be more lenient, i.e. choosing a SSNR of 0.25 (corresponding to a FSC of 0.2) instead of a SSNR of 0.167 for 3D in addition to accepting a window radius when 90 % of measurements (rather than the 95 % for 3D) passed. This design was chosen in order to limit the window size for our 2D measurements **(Suppl. Figure 2b)**.

### Band-pass filtering

For the employed band-pass filters in the real-space correlation approach, we used a hyperbolic tangent. As we wanted to employ a consistent band-pass filter independent of map size, and we did not choose a fixed number of pixels in Fourier space. For the bandwidth, we used 10 % of Nyquist centered at the current spatial frequency of interest combined with a two-sided symmetric falloff width of 1.5 %. Considerations were to choose the band width wide enough for reliable results, while still being small enough to be precise in Fourier space. Moreover, we padded input half-maps to the closest dimension for efficient fast Fourier transformation (FFT) calculations, and small maps received zero padding up to a minimum of 160 pixels in each dimension in 3D and 300 pixels in 2D.

### Threshold determination and reference distribution creation

For *p*-value determination, we require a reference distribution created via correlation calculations between one of the half-maps and its permuted counterpart. For refined maps (derived from single-particle cryo-EM or subtomogram averaging), we create this permuted half-map by random permutation in real space, preserving only the intensity distribution, as we do not expect systematic artifacts to be present due to the averaging of many different maps. However, for maps derived from single recordings, such as for micrographs, tilt-series and tomograms, systematic artifacts may arise from both acquisition and processing. One particularly relevant case is the missing wedge in tomograms, which creates anisotropic resolution and therefore causes correlated noise patterns across half-maps, artificially inflating correlations. For accurate statistics, it is necessary that the permuted half-map with which we derive our reference distribution also contains these artifacts. To capture these changes, we use phase-permutation instead of real-space permutation (**Suppl. Figure 1**), preserving the power spectrum while disrupting spatial correlations. Consequently, both the original and permuted maps share the same spectral properties and systematic biases, ensuring that our correlation measurements are tested against a realistic null distribution.

To keep the statistical properties of the permuted half-map truthful to the experimental input half-maps, it also needs to be subjected to the same band-pass filter as the input half-maps. Then, 10,000 correlation measurements between the band-pass-filtered permuted half-map and one of the original half-maps are conducted to generate the reference distribution, using the same window radius as for the actual half-maps, but at random positions. One potential limitation with this approach for reference distribution creation arises for situations with large windows and small maps, where the number of unique non-overlapping measurement locations is small, e.g. in the case of micrographs: repeatedly using the same or overlapping windows for conducting correlation measurements narrows the reference distribution. One simple solution we implemented for such cases is the creation of multiple permuted maps. Additionally, it is noteworthy that Fourier-space dependencies arise independently of real-space location. Therefore, the real-space window locations between both maps do not need to be identical. This random pairing dramatically increases the number of possible combinations. Subsequently, we determine the *p-*value for each local measurement between the experimental half-maps based on the obtained reference distribution. We iterate through every shell of interest for the procedure. In the interest of computational efficiency, for input 3D maps, we use every second voxel as a measurement location, in order to still have overlapping windows for even the highest resolution shells. For 2D maps, every fifth pixel is sufficient, as the windows used are much larger. We also offer a fast mode intended for large maps or batch processing of large datasets. Here, only every third voxel is used for 3D, and, both for 2D and 3D measurements, the number of investigated shells is halved.

After iterating though every resolution shell, we end up with a new map, consisting of a list of shell-derived *p-*values at each location. As established before (Beckers and Sachse, 2020), for each location, we correct for multiple testing converting *p*-values into *q*-values (Benjamini and Yekutieli, 2001). Finally, a threshold of *q*=0.01 is used for the identification of significant signal. We use the conventional way of threshold determination: we iterate through shell-derived *q*-values list, starting with the lowest-frequency shell, in order to identify the highest resolution shell containing significant signal, stopping when the first *q-*value loses significance. For tomograms, tilt-series and micrographs, we instead use a cutoff of *q*=0.05, and, additionally, require two subsequent shells to be above the threshold. The cutoff is then chosen as the first of these two shells. This modulation is useful as for noisier maps, the risk of prematurely assigning low resolution is much higher. Additionally, this lowers the risk of loss of significance following the first CTF zero-transition, which leads to unwarranted lower *p*-values (**Figure 3 a-b**).

### Median resolution estimation

We also established a way to derive a global median resolution estimate we termed ‘median resolution’ from RESOLVE’s local measurements for micrographs, tilt-series and tomograms. We base this approach on a procedure of frequency-dependent *p*-value determination: For each frequency shell, we determine the median *p*-value (from the full set of *p*-values from all measurement locations within the map) and from this simple 1D list, we derive the *q*-values via multiple testing correction. This correction allows us to determine a resolution the same way as for single pixels or voxels in the above described practice for micrographs, tilt-series and tomograms: employing a cutoff value of *q*=0.05 to the per-shell *q*-values results in a single-resolution estimate. For better comparability across datasets, by default RESOLVE focuses on signal regions only (regions where our measurements passed the lowest frequency shell).

For more stringent comparisons, we also allow the user to supply an input mask, defining the signal regions for the median resolution estimate, and additionally an option to use the mean instead of the median. As tilt-series present a set of processed 2D images, we adjust the procedure slightly: Instead of taking just the median *p-*value per frequency, we first take the median *p-*value for each 2D tilt image, and then the mean *p-*value across all tilts to derive a single *p-*value per frequency, resulting in a simple 1D *q*-value list after multiple testing correction, which we use for cutoff determination as described previously, again resulting in a single-resolution estimate.

For *in situ* tomograms, we expect variability in resolution dependent on *z*-plane. For that reason, similar as for tilt-series, we take the median *p-*value for each *z*-slice, and then the mean *p-*value across all *z*-slices to derive a single *p-*value per frequency. This is, as for tilt-series, followed by multiple testing correction and cutoff determination resulting in a single-resolution estimate. Generally, for tomograms, tilt-series and micrographs, RESOLVE automatically produces a frequency dependent median *q-*value plot from which the median resolution is derived. Additionally, for tilt-series and tomograms, RESOLVE also produces a *p*-value plot, displaying tilt or z-slice-dependent *p*-value behavior (**Fig. 4a, middle, heatmap**). Together with the optional user-defined input mask, allowing to focus these measurements on regions of interest, this setup allows for maximal flexibility and sensitivity in detecting small quality changes.

### Single-particle image processing

For the TMV micrographs (EMPIAR-10305), frames were motion corrected and split into half-frames with MotionCor3 (Zheng et al., 2017) using a pixel size of 2.925 Å for resolution measurements. For CTF estimation, we used CTFFIND4 (Rohou and Grigorieff, 2015) and for the single-particle workflow with EMPIAR-10524, we used RELION-5 (Burt et al., 2024). Micrograph half-sets were reconstructed from raw frames with MotionCor3 (Zheng et al., 2017). After motion correction, half-micrographs were binned with IMOD (Mastronarde, 2024) by a factor of 2 to a pixel size of 2.57 Å. RESOLVE was applied to the binned, non CTF-corrected micrograph half-sets. For refinements with preselected particles by RESOLVE’s micrograph resolution estimates, particles on micrographs with carbon film were excluded. For estimating particle quality on a micrograph level, a custom script was used to associate star file coordinates and micrograph resolution. Particle quality was estimated by averaging all local RESOLVE resolution estimates within an approximate 10 nm diameter box around the particle location. Micrographs were visualized with IMOD (Mastronarde, 2005).

### Tomography image processing

For the tomography dataset EMPIAR-11306, we employed the tomography pipeline of RELION-5 (Burt et al., 2024), including half-tilt-series and half-tomogram reconstructions. Half-tilts were stacked to produce a half-tilt-series and binned using IMOD (Mastronarde, 2024). Resolution measurements were conducted for the tilt-series with a final pixel size of 11.1 Å after binning. Half-tomograms were reconstructed with a pixel size of 12 Å. We used CTFFIND4 for CTF estimation (Rohou and Grigorieff, 2015). For tilt-series alignment, we used AreTomo (Zheng et al., 2022). For estimating particle quality on a tomogram level, a custom python script was used to associate star file coordinates and tomogram resolution as measured by RESOLVE. Resolution estimates on tomograms were conducted with the RELION-5 reconstructed tomogram half-maps. Particle quality was estimated by averaging all local RESOLVE resolution estimates within an approximate 20 nm diameter box around the particle location. For local ribosomes resolution estimated by RESOLVE as portrayed in **Suppl. Figure 5** and **Suppl. Figure 6**, the particle set after the first 3D classification was used (∼ 27,500 ribosomes). Lamellae were modelled for 106 tomograms. For better lamella visibility, tomograms were denoised with cryoCARE (Buchholz et al., 2019). Tomograms where the lamella surface was difficult to identify were excluded. For the remaining 106 tomograms, top and bottom lamella surfaces were marked manually for multiple y-slices throughout the tomograms, so that the surfaces could be interpolated (**Suppl. Figure 4**). Distances to surfaces were determined for every second voxel, using the closest surface coordinate as reference. Local lamella thickness was estimated as the distance between closest top surface coordinate and closest bottom surface coordinate. The tomogram visualizations were created with either IMOD or Chimera (Pettersen et al., 2004).

### Software and hardware specifications

We provide RESOLVE as a standalone tool, including four different modes of processing: 1) Refined maps, referring to single-particle and subtomogram average maps 2) 2D micrographs 3) tilt-series 4) tomograms. RESOLVE comes with a graphical user interface and can also be run in command-line mode to support batch processing tasks. The code and tutorial is shared at https://github.com/DavidKart/RESOLVE. On our workstation equipped with a single NVIDIA GeForce RTX 3060 and an AMD Ryzen 9 7900X 12-core processor, RESOLVE required approximately 5 minutes to process a large input tomogram (560 MB), with processing time reduced to 2 minutes when fast mode was enabled (utilizing lower sampling frequencies in both Fourier and real space). This calculation consumed 14 GB of RAM and 3 GB of GPU memory. For large input maps, distributing the workload across multiple GPUs can accelerate processing, though we recommend using no more than 2 GPUs to maintain efficiency. For single micrographs (2.7 MB), processing took only 12 seconds in standard mode and 7 seconds in fast mode. Notably, GPU acceleration is not recommended for such small maps, as it reduces performance.

## Supplementary Figures

**Supplementary Figure 1:**
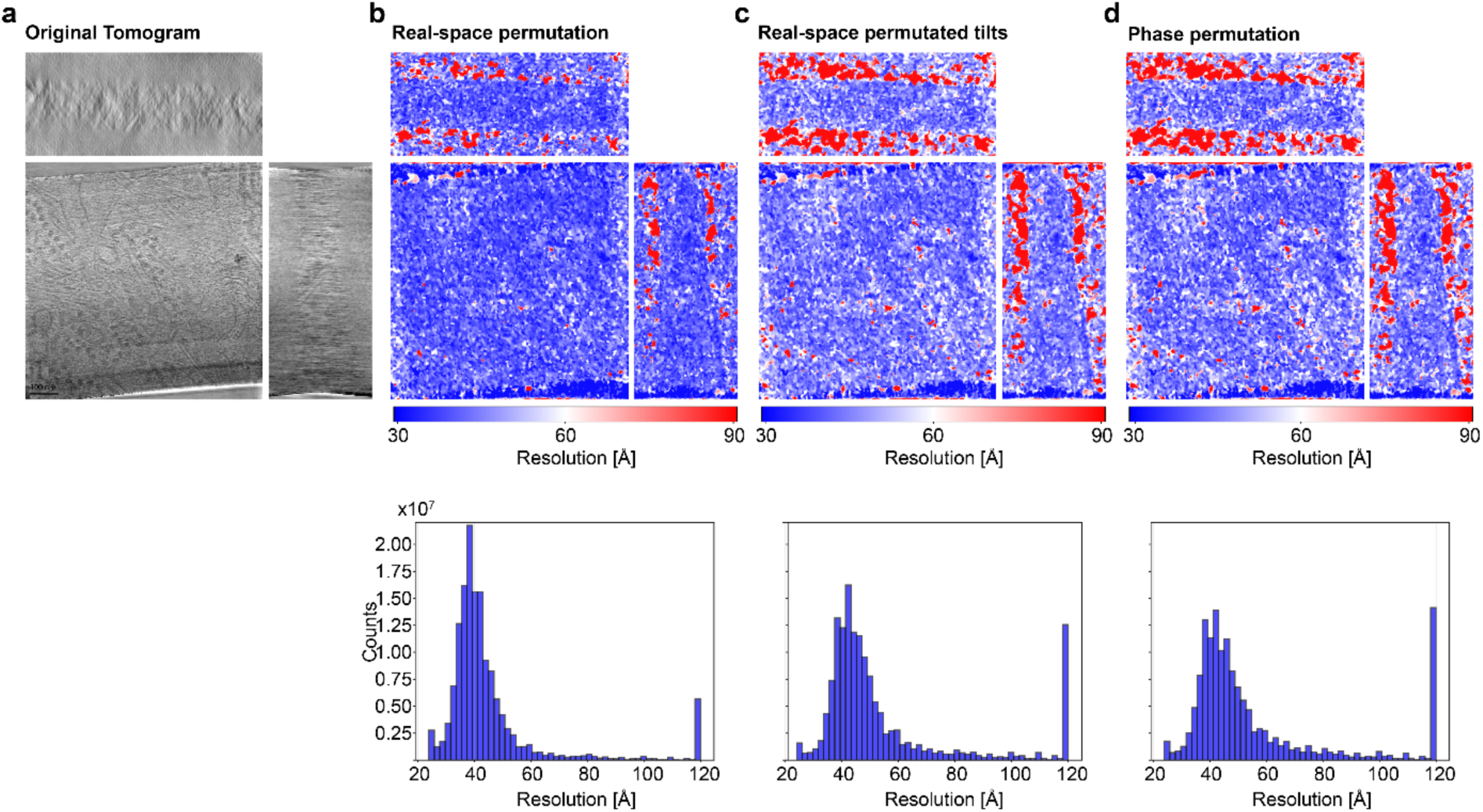
Different permutation methods for tomogram reconstruction and effect on resolution estimates. Top row: central slices of the three axes. Bottom row: Histograms of resolutions of the complete tomograms. From left to right: a) Original, denoised tomogram. b) Resolution measurement using real-space permutation for reference distribution creation. c) Resolution measurement conducted by using a tomogram as reference distribution created by real-space permutation of the single tilt images before reconstruction, ensuring any potentially biases introduced via tomogram reconstruction are included in the reference distribution. d) Resolution measurement using phase-permutation for reference distribution creation.

**Supplementary Figure 2:**
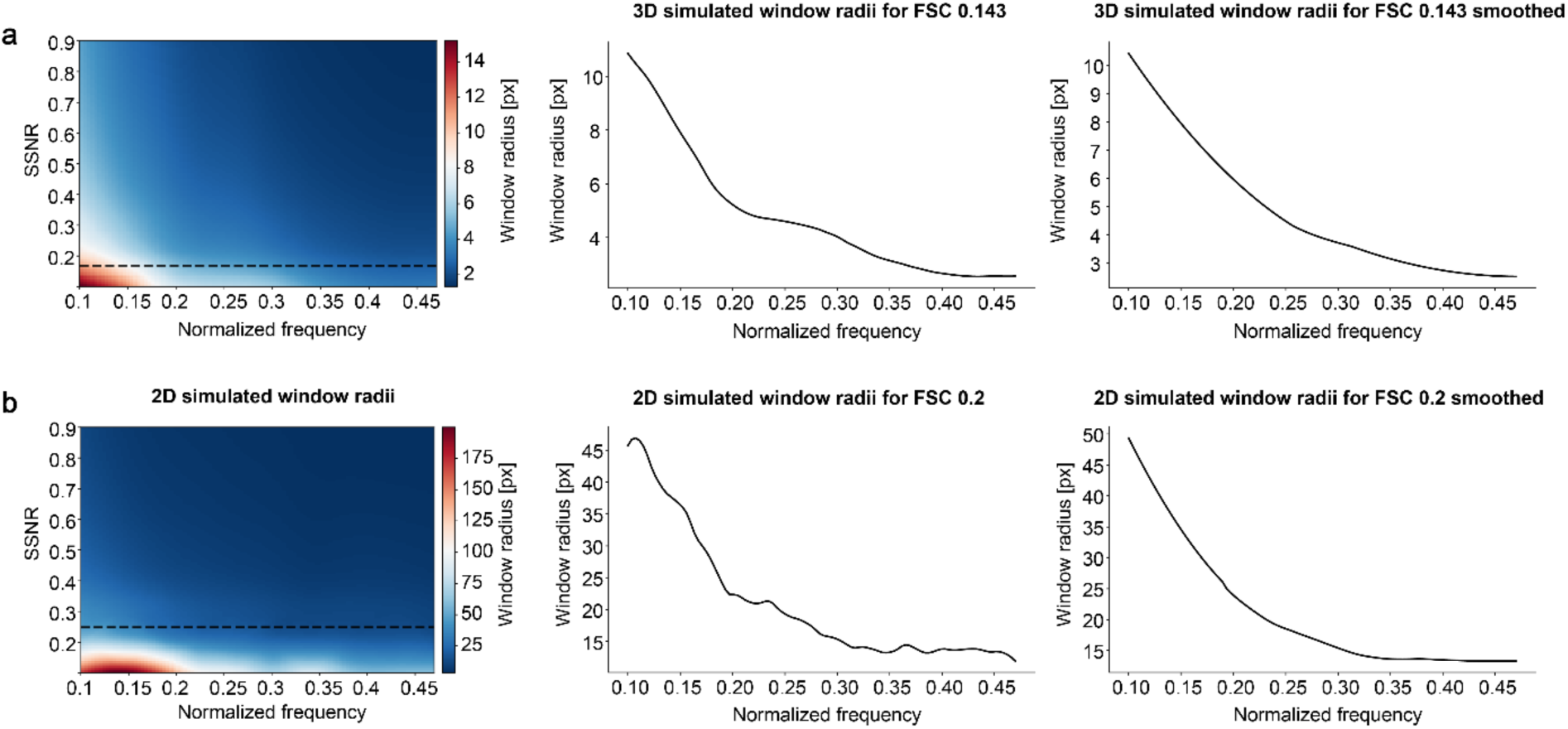
Window radii simulations for three-dimensional (3D) and two-dimensional (2D) data types. a) Window radii determined from simulation for 3D measurements. Left: Table with the determined frequency dependent window radii for different spectral signal-to-noise (SSNR) levels. SSNR 0.167 (FSC 0.143) is highlighted with a black dotted line. Middle: Extracted original curve for SSNR 0.167. Right: Smoothed curve via a Savitzky-Golay filter for SSNR 0.167. b) Window radii determined from simulation for 2D measurements. Left: Table with the determined frequency dependent window radii for different SSNR levels. SSNR 0.25 (FSC 0.2) is highlighted with a black dotted line. Middle: Extracted original curve for SSNR 0.25. Right: Smoothened curve via a Savitzky-Golay filter for SSNR 0.25.

**Supplementary Figure 3.**
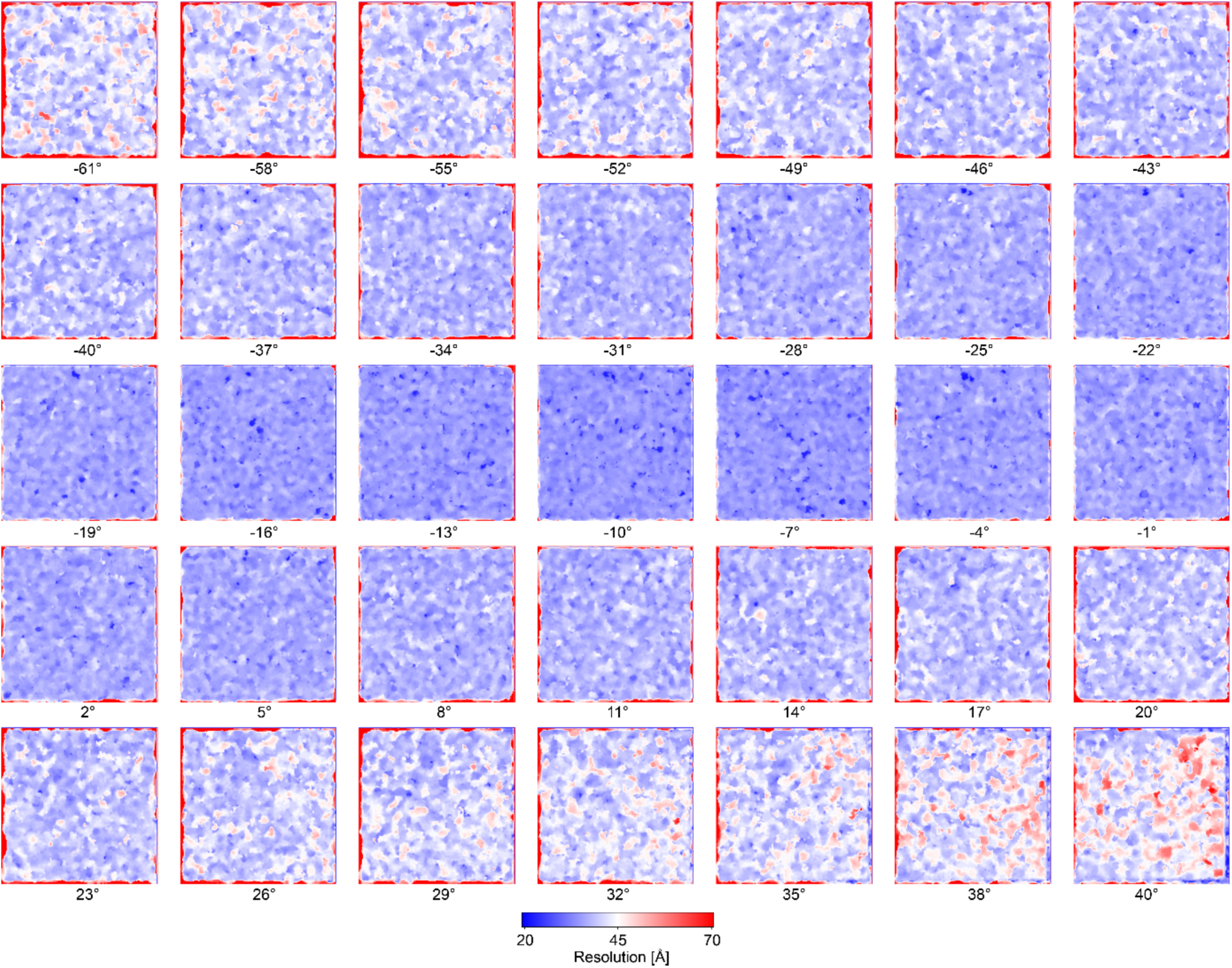
Resolution maps (color bar below from 20 to 70 Å) for full tilt series of 35 tilted micrographs from -61 to 40°.

**Supplementary Figure 4:**
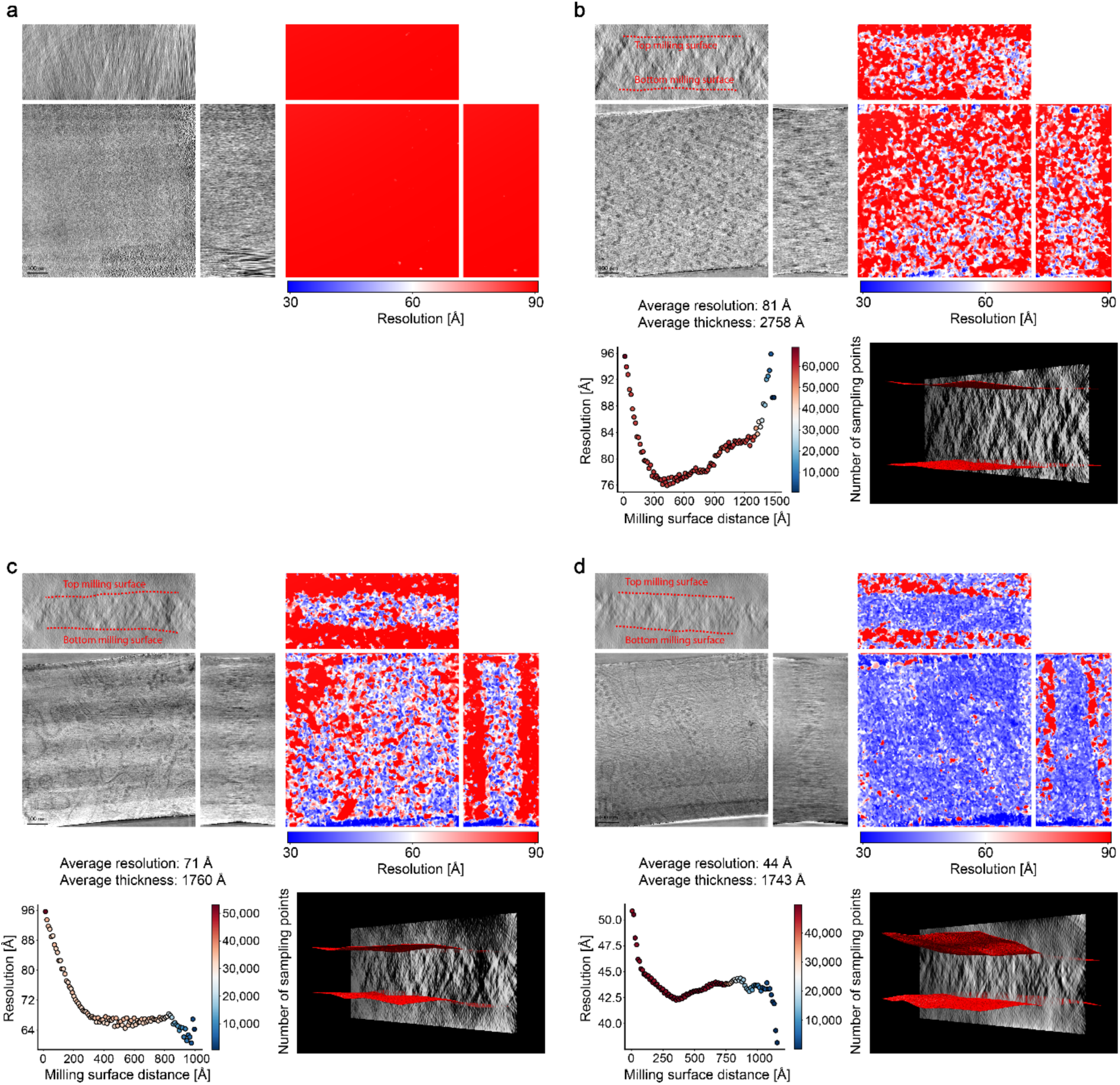
Lamella model and resolution estimates of four illustrative tomograms. a) Exemplary empty tomogram. Representative tomograms of b) a thick lamella, c) a thin lamella and d) another thin lamella. Central slices through the three axes visualized as grey-scale intensities (top left) and the corresponding resolution mapping (top right). Bottom left shows averaged resolution values dependent on distance to closest milling surface for the corresponding tomogram. Only voxels within the modeled lamella are included in the measurement. Bottom right shows lamella model though central xz-slice.

**Supplementary Figure 5:**
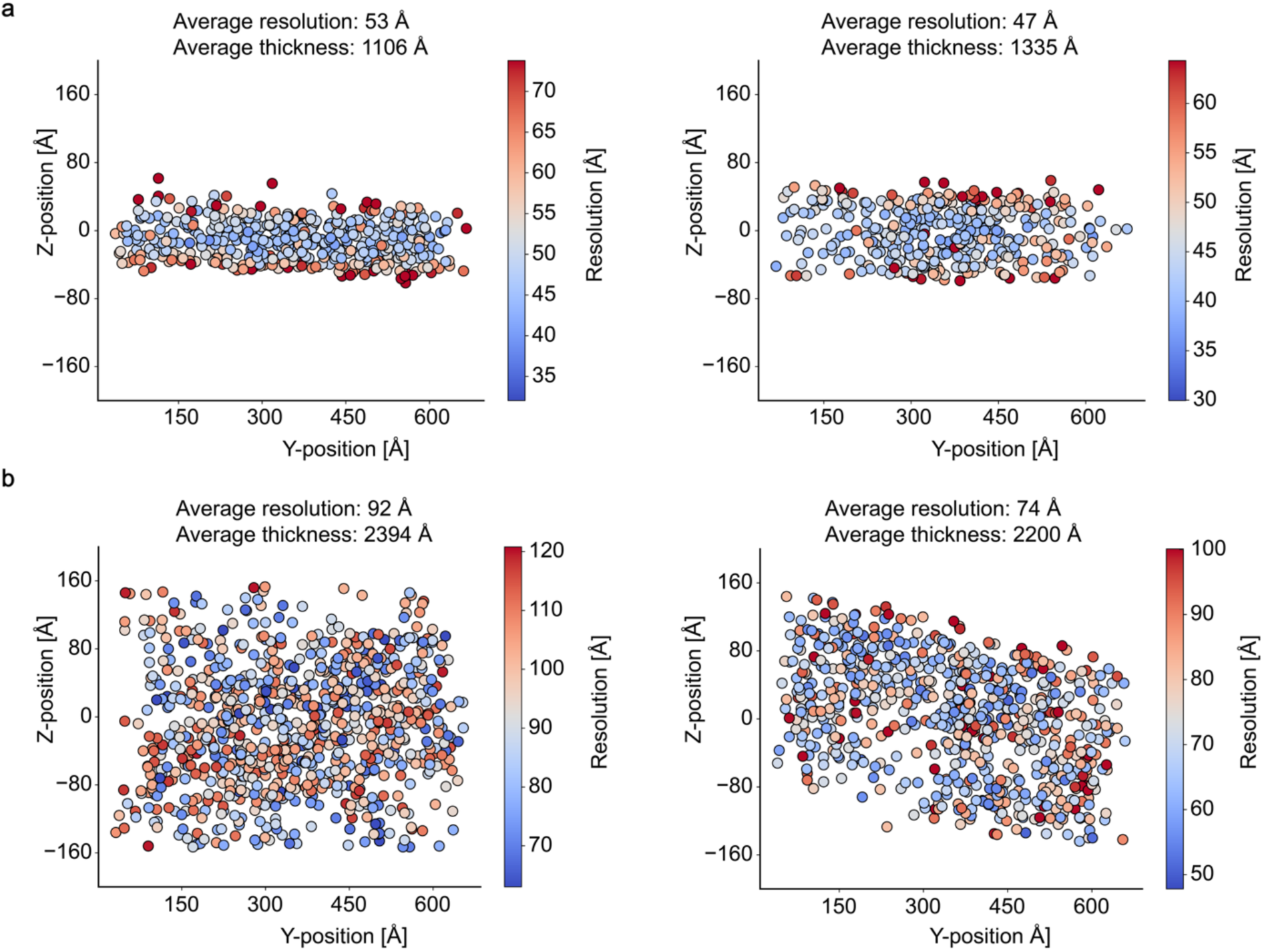
Local resolution estimates based on the Z-position of ribosomes within lamellae. RESOLVE’s local resolution estimates based on ribosomes mapped on Y and Z positions in the tomogram with color bar on the right: a) Local resolution estimates of two representative thin lamellae (left and right). b) Local resolution estimates of two representative thick lamellae (left and right).

**Supplementary Figure 6:**
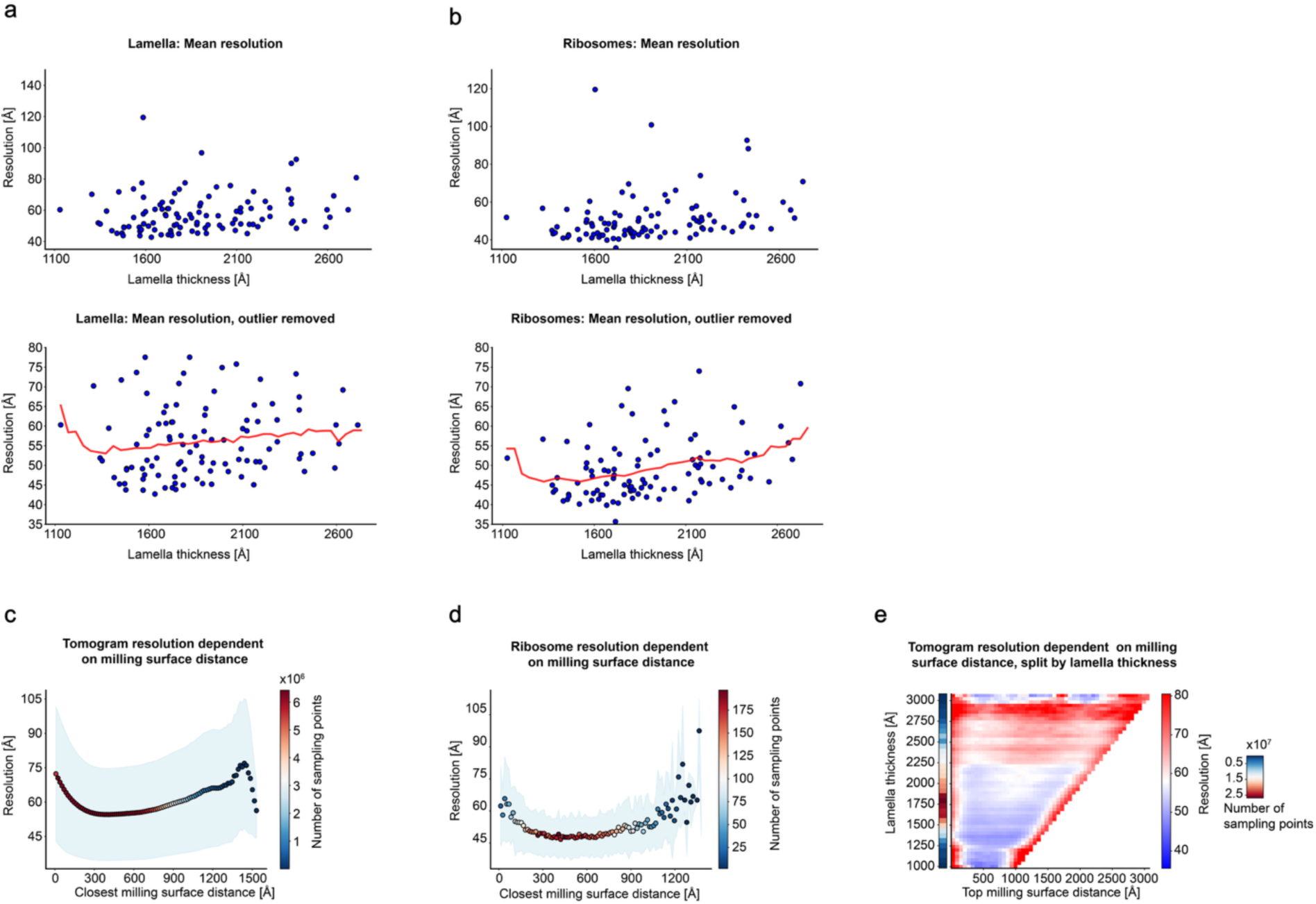
Local resolution vs. thickness and resolution distribution within tomograms. a) Scatter plot of average resolution per tomogram within the lamella vs. lamella thickness. Top: full dataset (106 tomograms). Bottom: Outliers > 80 Å removed, moving average as red line (window size 400 Å). b) Scatter plot of average resolution of ribosomes (averaged per tomogram) within the lamella vs. Lamella thickness. Top: full dataset (102 tomograms). Bottom: Outliers > 80 Å removed, moving average as red line (window size 400 Å). c) Average resolution vs. distance to closest milling surface for 106 tomograms. Blue background coloring portrays standard deviation. d) Average resolution vs. distance to closest milling surface focused on ribosomes. Blue background coloring portrays standard deviation. e) Lamella thickness vs. distance to top milling surface, resolution and particle numbers are color coded. Full set of 106 tomograms.

